# Shear forces drive precise patterning of hair cells in the mammalian inner ear

**DOI:** 10.1101/707422

**Authors:** Roie Cohen, Liat Amir-Zilberstein, Micha Hersch, Shiran Woland, Shahar Taiber, Fumio Matsuzaki, Sven Bergmann, Karen B. Avraham, David Sprinzak

## Abstract

Precise cellular organizations are required for the function of many organs and tissues. It is often unclear, however, how such precise patterns emerge during development. The mammalian hearing organ, the organ of Corti, consists of a remarkably organized pattern of four rows of hair cells (HCs) interspersed by non-sensory supporting cells (SCs). This checkerboard-like pattern of HCs and SCs emerges from a disordered epithelium over several days, yet the transition to an ordered cellular pattern is not well understood. Using a combination of quantitative morphological analysis and time-lapse imaging of mouse cochlear explants, we show here that patterning of the organ of Corti involves dynamic reorganizations that include lateral shear motion, cell intercalations, and delaminations. A mathematical model, where tissue morphology is described in terms of the mechanical forces that act on cells and cellular junctions, suggests that global shear on HCs and local repulsion between HCs are sufficient to drive the tissue into the final checkerboard-like pattern. Our findings suggest that precise patterns can emerge during development from reorganization processes, driven by a combination of global and local forces in a process analogous to shear-induced crystallization.

## Main text

The mature organ of Corti (OoC) is a strip of epithelial cells that extends along the base-to-apex axis within the cochlear duct (Fig. 1a,b) and includes three outer hair cell (OHC) rows and one inner hair cell (IHC) row, regularly interspersed by the non-sensory SCs. The OHC region is flanked by the Hensen cells on its lateral side and a single row of cuboidal-shaped inner pillar cells (PC) that separate it from the IHCs on its medial side. Studies in mice showed that the differentiation into HCs and SCs initiates from a disordered undifferentiated layer of cells at around embryonic day 14 (E14), and involves a lateral inhibition process mediated by Notch signaling^1–3^. At the same time, the OoC exhibits significant morphological changes that include convergent extension^4–6^ and cell movements^7^. No significant cell division or cell death events are observed within the OoC during this process^4,8^.

**Figure 1.**
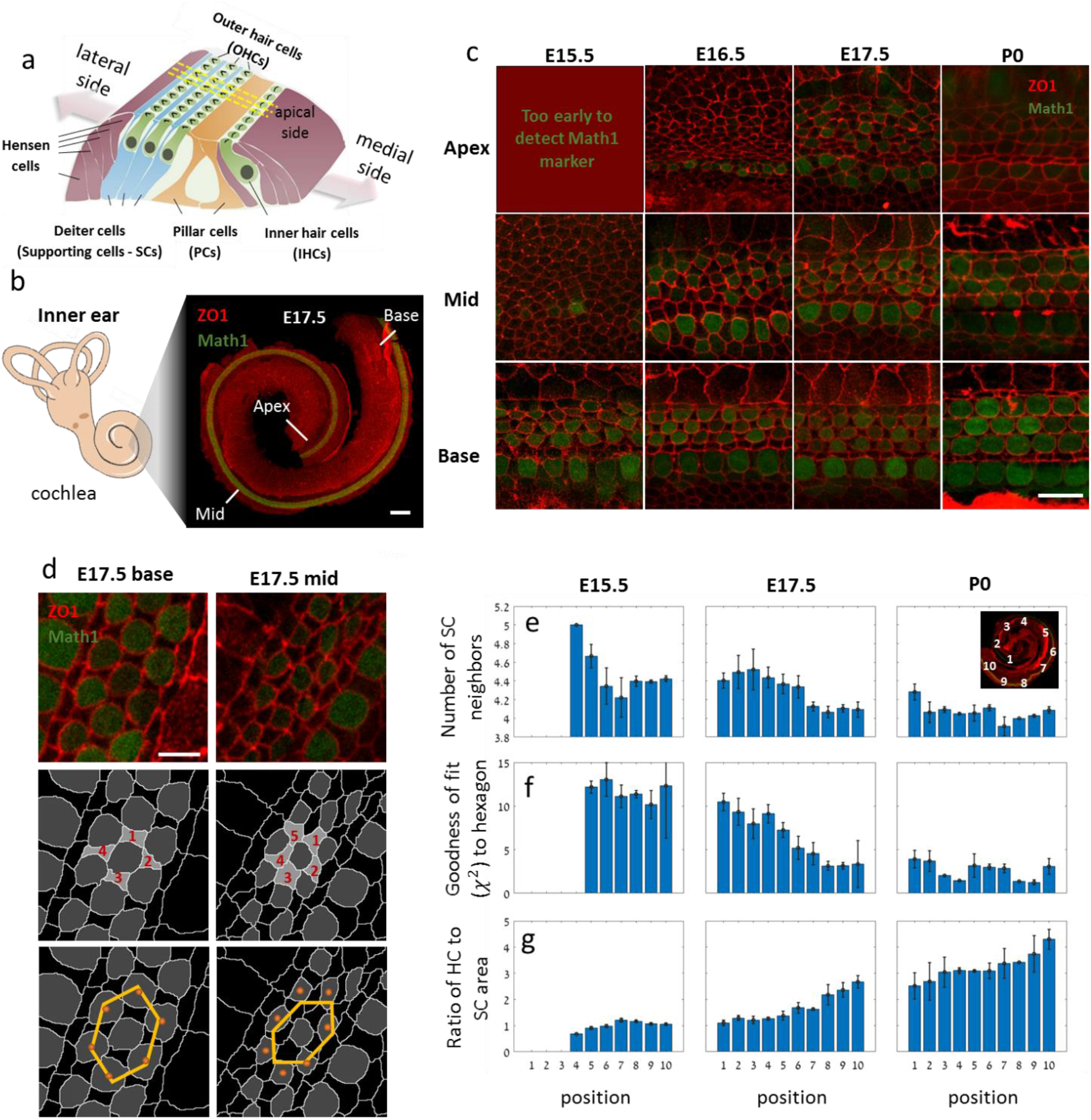
Hair cells gradually reorganize into a compact checkerboard-like pattern. **(a)** Schematic of the OoC. **(b)** Left: schematic of the mammalian inner ear. Right: Confocal image of the cochlea of a E17.5 mouse embryo marking the base, mid and apex regions. Cochleae are taken from transgenic mice expressing Math1-GFP (green) and are immunostained with α- ZO1 (red). Scale bar: 100*μm*. **(c)** Representative images at different developmental times and different positions along the base-to-apex axis. Rows and columns correspond to different positions along the cochlear axis and different development times, respectively, as indicated. Scale bar: 10*μm*. **(d)** Schematic of the definition of two order parameters: (i) The number of SC neighbors of each HC in OHC_2_ (middle row) and (ii) goodness of fit to a symmetric hexagon (bottom row). Symmetric hexagon is defined as a regular hexagon with one axis multiplied by a scaling factor. Scale bar: 5*μm*. **(e-g)** Morphological and order parameters in different regions of the cochlea from apex to base (defined in inset) and at different developmental times (columns). Rows correspond to number of SCs neighbors (e), goodness of fit to a symmetric hexagon (f), and ratio of HC to SC surface area (g). The data are averaged over n=3,4,3 cochleae at E15.5, E17.5, P0, respectively. Schematic in (a) is modified with permission from Dror and Avraham^11^.

To determine the processes that drive the transition from a disordered undifferentiated tissue to an ordered pattern of HCs and SCs we first quantitatively analyzed the morphological changes that occur in the OoC in space and time. We utilized the fact that the organization of the OoC exhibits a developmental gradient along the base-to-apex axis^1^, allowing continuous analysis of the patterning progression in fixed samples. To this end, we imaged fixed cochleae at different developmental stages (E15.5 to postnatal day 0 (P0)) taken from transgenic mice expressing Math1-GFP^9^, which is an early marker of HC differentiation. The cochleae were additionally immunostained with an antibody against zonula occludens-1 (ZO1), a known marker of tight junctions^10^, delineating cell boundaries at the apical side of the tissue (Fig. 1a). Images of full cochleae were reconstructed by tiling multiple high-resolution confocal images taken at different Z-planes. Images were then segmented and analyzed using a custom-built code (Fig. S1a) allowing extraction of morphological parameters (e.g. cell size, number of neighbors) and cell identity (HC or SC).

The analysis both along the base-to-apex axis and at different time points demonstrates the gradual differentiation and organization of the OoC in both space and time (Fig. 1c). To obtain a quantitative measure of the level of organization we defined two local order parameters. The first parameter is the number of SC neighbors for each HC in the OHC region. In the final pattern, each OHC from the middle HC row (OHC_2_) has exactly four SC neighbors, while at earlier stages this number is often higher (Fig. 1d, middle row). Analysis of cochlea from E15.5, E17.5, and P0 shows that the average number of SC neighbors decreases as development progresses (both in space and time, Fig. 1e).

The second order parameter defines a measure for the hexagonal packing of HCs. In a fully ordered OoC, each HC from OHC_2_ is at the center of a symmetric hexagon (a regular hexagon squeezed in one axis) formed by its closest HC neighbors (Fig. 1d, bottom row). To obtain a measure of the hexagonal order, for each HC in OHC_2_, we look at the polygon formed by the centers of its neighboring HCs, and find a symmetric hexagon that best fits that polygon (yellow hexagons in Fig. 1d). We define the goodness of fit to hexagon (*χ*^2^) as our measure of hexagonal order. This parameter is lower in value at more advanced developmental stages and higher at less advanced stages indicating the gradual organization of the HCs into a hexagonal pattern (Fig. 1f). Consistent with earlier works^12^, we also find that the apical surface areas of the HCs increase with developmental stage, while the corresponding areas of the SCs slightly decrease (Fig. 1g, Fig S1b). Overall, this analysis shows that gradual transition into an organized pattern involves a decrease in the number of SC neighbors of each HC, an increase in the hexagonal order of HCs, and an increase in the relative areas of HCs with respect to that of SCs.

To understand the dynamics underlying the morphological changes that take place during the development of the OoC, we developed an assay for live imaging of inner ear explants. In this assay, cochlear explants from transgenic mice expressing the boundary marker ZO1-EGFP^13^ were imaged at high resolution using a confocal microscope equipped with an Airyscan detector, for up to 48 hours. Movies of cochlear explants were performed on cochlea extracted at E15.5, E17.5, and P0.

Remarkable reorganization processes were observed at earlier developmental stages. Imaging of E15.5 explants, showed a significant shear motion of the HCs in the direction of cochlear extension (Fig. 2a-b, Supplementary Video 1). In particular, we observed shear motion between outer and inner HCs, where the outer HCs slide in an opposite direction relative to inner HCs. Analysis of the relative displacement of cell layers using an image registration algorithm, showed that regions farther away from the pillar cell layer (dashed line in Fig. 2a) exhibited larger displacement than regions closer to the pillar cell layer (Fig. 2c top, Fig. S2c left). Furthermore, the displacement in the medial-lateral axis (perpendicular to the base-apex axis) showed the tendency of the tissue to converge towards the pillar cell layer (Fig. 2c bottom, Fig. S2c right). We also note that the Hensen cells, medial to the OHCs, are highly dynamic in their shape and exhibit directional sliding movement with respect to the OHC layer even at more advanced developmental stages (Fig. S2d, Supplementary Video 2). Unlike the cells in the OoC, these cells continue to divide even at much later developmental stages^8^. Overall, these observations suggest that OHCs undergo shear motion, potentially driven by dynamic movement and cell division of Hensen cells.

**Figure 2.**
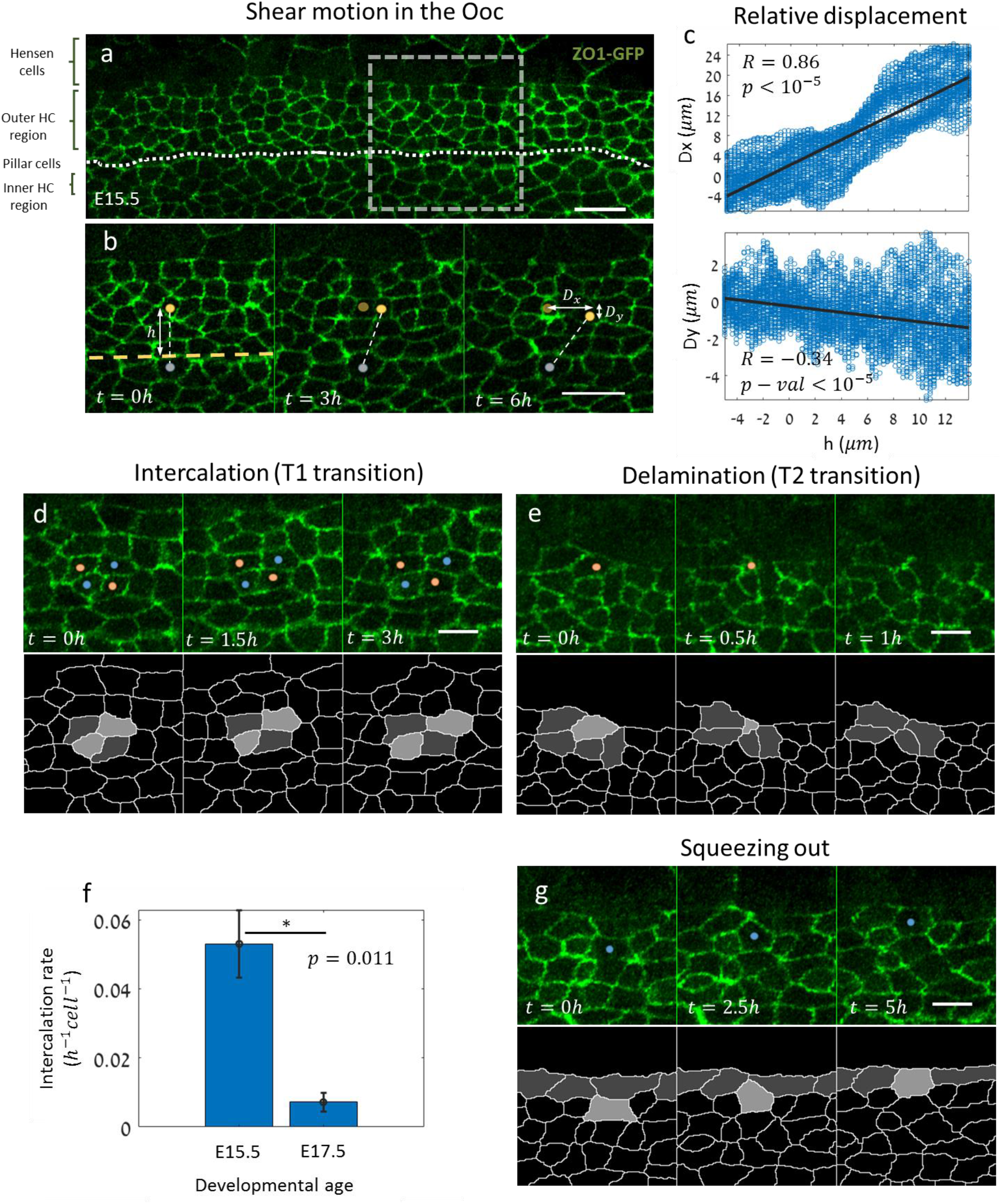
Shear motion and local morphological transitions drive organization in the organ of Corti. **(a)** A filmstrip from the mid-apex region of a cochlear explant of a ZO1-EGFP mouse at E15.5. Top: Image of a section of the OoC showing the different regions along the medial-lateral axis. Dotted line marks the pillar cell (PC) row. **(b)** A filmstrip from a time-lapse movie (dashed square region in (a)) showing shear motion in the OHC region. A connecting line between two cells in the OHC region (yellow dot) and IHC region (gray dot) highlights the relative motion between the two marked cells. *h* is the initial distance from the PC row and *D*_*x*_, *D*_*y*_ are the total displacements in the *x* and *y*; directions at the end of the movie compared to the original position (orange dot). Scale bars: 10*μm*. Movie shown in Supplementary Video 1. **(c)** Total pixel displacement in the filmstrip shown in (a) as a function of the initial distance from the PCs. Pixel displacement is calculated using an image registration algorithm (methods). Black lines represent linear regressions for the horizontal (top) and perpendicular (bottom) movements. The correlation coefficients (R) and corresponding p-values are as denoted. **(d)** A filmstrip from the same OoC region in (a) showing an intercalation process (T1 transition) between the cell pair marked in red dots and the cell pair marked in blue dots. Bottom row shows a segmented version of the intercalation process. Movie shown in Supplementary Video 3. **(e)** A filmstrip from the same OoC region in (a) showing a delamination process (T2 transition), where the cell marked with red dot decreases in surface area and eventually vanishes from the apical surface. Bottom row presents a segmented version of the transition. Movie shown in Supplementary Video 4. **(f)** Rate of intercalations in the OoC (# of intercalations per hour per cell) at E15.5 and E17.5. Data obtained from n=3 movies. Error bars represent S.E.M. **(g)** A filmstrip from the same OoC region in (a) showing an event where a cell (blue dot) is “squeezed out” towards the top border of the OHC region. Bottom row presents a segmented version of the process. Movie shown in Supplementary Video 7. Scale bars for (d,e,g): 5*μm*.

Previous studies of developing epithelial tissues showed that global morphological changes often involve local morphological transitions including cellular intercalations (T1 transitions) and delaminations (T2 transitions) ^14,15^. Analysis of our movies indeed showed both intercalations and delaminations both at E15.5 (Fig. 2d-e, Supplementary Videos 3,4) and at E17.5 (Fig. S2c-d, Supplementary Video 5,6). We find that the number of intercalations observed is significantly higher at earlier developmental stages (E15.5) compared to later developmental stages (E17.5) (Fig. 2f), and inversely correlated with the degree of tissue organization.

A specific type of reorganization observed in our movies, is a situation where a cell, which is initially located within the middle of the OHC region, is ‘squeezed’ laterally to the row of cells that borders the Hensen cells (Fig. 2g, Supplementary Video 7). Both lateral squeezing and cellular delaminations seem to be restricted to cells with SC morphology (more concave shaped). Both of these processes may contribute to the reduction in the number of SC neighbors of HCs shown in Fig. 1e.

To get an insight into the processes that lead to precise patterning of the OoC, we developed a mathematical model of the patterning process. As a first stage, we wanted to determine the minimal rules that can capture the transition from a disordered epithelial layer to an ordered checkerboard-like pattern, focusing on the OHC patterning (Fig. 3a). We chose to use a 2D vertex model, previously used to describe morphological transitions in growing epithelial tissues^15,16^. In 2D vertex models, different cells are attributed with different mechanical properties associated with the cell area, cell perimeter, and the cellular junctions, while the cellular configuration is determined by minimizing the overall mechanical energy of the system (Fig. S3a). For the model of the OoC, each cell is defined as a HC, SC, boundary cells (e.g. PCs and top boundary cells), or a general cell outside the OoC (Fig. 3a). In parallel to the minimization of the mechanical energy we introduced in the model intercalations and delaminations that facilitate morphological changes. Intercalations and delaminations initiate when the junction or cell area fall below some threshold length or threshold area, respectively. For simplicity, we use periodic boundary conditions at the edges of the cell lattice.

**Figure 3.**
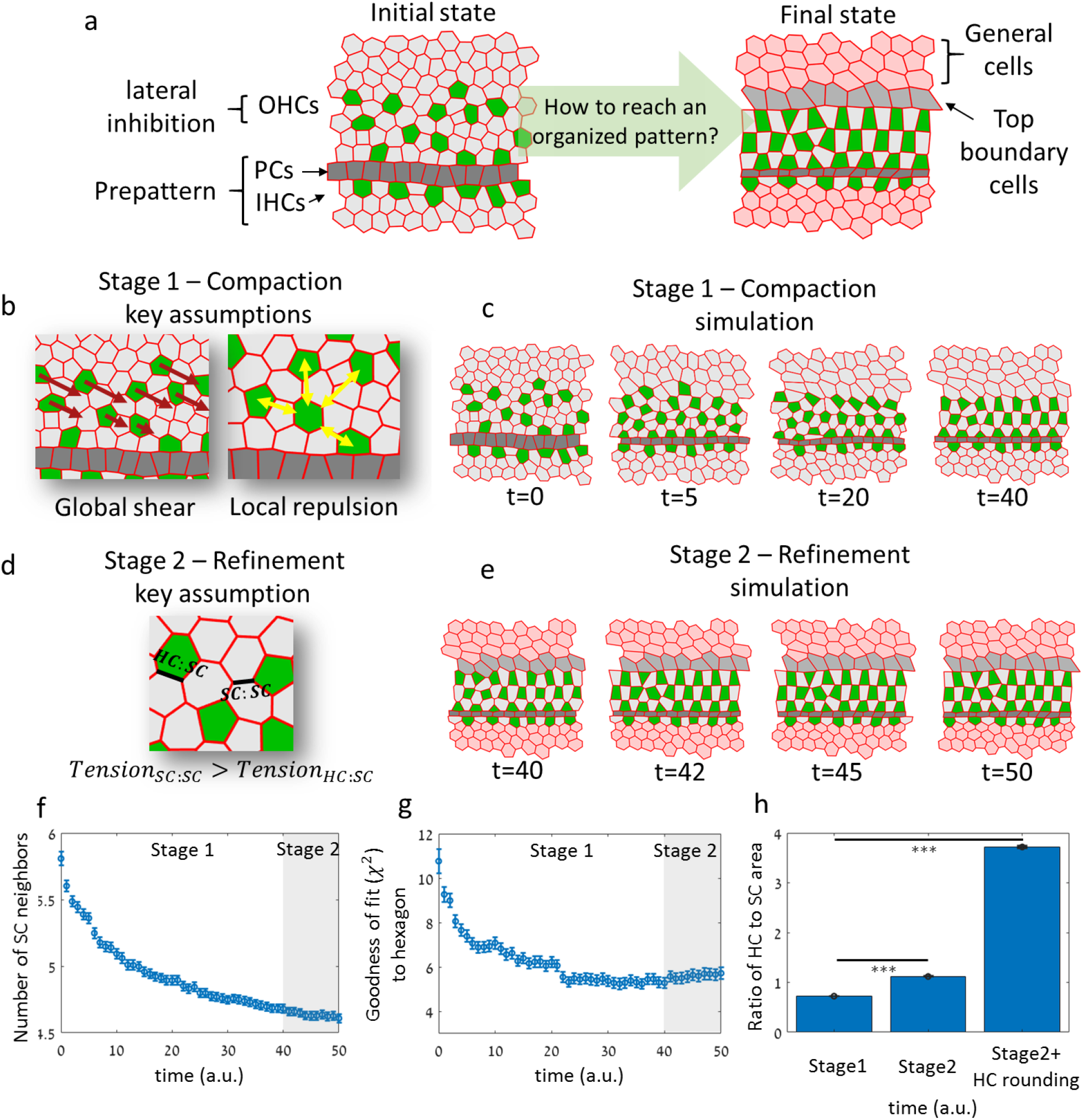
A mechanical model based on global shear and local repulsion explains precise patterning of the OHCs. **(a)** Schematic of the initial state of the model and the desired final state. The initial state for the simulation begins with the IHCs and PCs already pre-formed. A lateral inhibition mechanism is used to define a disordered pattern of HCs and SCs at a certain distance from the PCs. **(b)** Schematic of the key assumptions for the first stage of the simulation (compaction stage). HCs are subjected to global shear forces (red arrows) and local repulsion forces (yellow arrows). **(c)** A filmstrip of the first stage in the simulation. Global shear and local repulsion forces lead to the formation of a compact state of HCs. Movie shown in Supplementary Video 8. **(d)** Schematic of the key assumption for the second stage (refinement stage). The tension in SC:SC junctions is increased relative to that of HC:SC junctions. **(e)** A filmstrip of the second stage in the simulation. This stage begins at the end of the compaction stage. In addition to the key assumption in this stage, a top border, similar to the one between OHCs and PCs is introduced in these simulations. Movie shown in Supplementary Video 9. **(f-g)** Simulations capture the dynamics of the order parameters observed experimentally. Similar to the analysis in Fig. 1e-f, the number of SC neighbors decreases (f) and the HC organization became more hexagonal (g). **(h)** Simulations capture change in HCs and SCs areas. Bars correspond to the ratio of HC to SC areas at the end of stage 1, stage 2, and after artificial rounding of HCs (Fig. S3b). The data is averaged over n=50 simulations. Error bars in (f-h) indicate S.E.M. Full description of the simulations is provided in the Methods. Parameters used are provided in table S1.

Since we focus on the differentiation and patterning of the three OHC rows, and since the IHCs and PCs appear prior to the OHC differentiation (Fig. 1c), we began the simulations with IHCs and the adjacent rectangular-shaped PC row already formed (Fig. 3a, left). To simulate the initial alternating pattern of OHCs, we apply a lateral inhibition mechanism^17–19^, to a region extending laterally to the PC row. Lateral inhibition generates a disordered salt and pepper pattern in this region (Fig. 3a, left) and is further imposed throughout the simulations.

The goal of our model is to understand how reorganization processes can drive the initially disordered pattern in the OHC region into the final hexagonal pattern in the OoC. We have found that a key assumption required for achieving the hexagonal ordering is that there is an external force that compresses OHCs into a closed packed pattern. Based on the observation of shear motion of HCs in early developmental stages (Fig. 2a-b) and the evidence of convergent extension^6,7,12^, we modeled the external forces as diagonally oriented shear forces acting on the OHCs towards the PC row (Fig. 3b, left).

In addition to the global shear forces, we needed to take into account the observation that HCs rarely neighbor other HCs. To account for this behavior, we assume local repulsion forces between HCs (Fig. 3b, right). We note that both the global shear and the local repulsion forces operate on HCs and not SCs. This is justified by the fact that at the level of the nuclei, the HCs are physically separated from the SCs (e.g. the nuclei of SCs and HCs are on different planes, see Fig. 1a). Furthermore, unlike the SCs, the HCs are not anchored to the basement membrane and are therefore free to migrate^7^. In this context, the repulsion forces may therefore arise from HCs touching each other at the level of the nuclei. Since the HCs in our movies seem to be rounder than SCs and exhibit less changes in their shapes, we further assumed that the HCs in our simulations are more rigid (less compressible) and more contractile (tendency to maintain round shape) than the SCs (see methods). Simulations performed after applying global shear and local repulsions to the 2D vertex model capture the main features of the patterning process of OHCs (Fig. 3c, Supplementary Video 8). More specifically, it captures the transition into a hexagonal compact state of HCs and the decrease in the number of SC neighbors.

A second aspect we wanted to capture in the model was the increase in the area of HCs relative to that of the SCs (Fig. 1g). Given the observed decrease in length of junctions between two supporting cells (SC:SC junctions) compared to the length of the junctions between HCs and SCs (HC:SC junctions), we introduced a second stage in our simulation in which we increased the SC:SC tension (Fig 3d). At this stage we also introduced high tension at the top boundary to constrict the OHC region. Applying these additional assumptions resulted in an almost-perfect confined checkerboard-like pattern (Fig. 3e. Supplementary Video 9).

Analysis of the order parameters in our simulated data showed that the number of SC neighbors and the goodness of fit to hexagon for OHCs, decrease with simulation time (Fig. 3f-g) similar to the experimental behavior (Fig. 1e-f). The simulations also show that the reduction in the number of SC neighbors is due to SCs delaminating or squeezing out toward the lateral layer. Although the second stage did not improve the order parameters, it was essential to obtain the observed final structure of the OoC and captures the relative change in HC and SC area (Fig. 3h). We note that, for simplification purposes the first and second stages are modeled sequentially although these process may actually overlap.

The proposed model does fail to capture some aspects of the final pattern of the OoC. In particular, the HCs in the model adopt a rectangular shape rather than the round shape observed. This is due to the fact that in 2D vertex models, cell shapes are restricted to polygons with straight boundaries. If we artificially introduce cell rounding to the HCs at the simulation endpoint (Fig. S3b), we observe an even higher similarity to the actual HC pattern and seem to capture the observed ratio between HCs and SCs areas (right bar in Fig. 3h). Physically, HC rounding may occur due to the effect of higher internal pressure in HCs^20^. We also note that our model does not achieve perfect hexagonal patterning as some defects are still observed. Again, additional assumptions may be required for the final tuning of the pattern.

We also wanted to test if alternative models, which do not require global forces, can also capture the transition into hexagonal patterning. A potential mechanism that can lead to organized patterning is preferential adhesion of HCs to SCs over adhesion of SCs to SCs. This is equivalent to assuming only stage 2 occurs in our simulations. Such a model was previously used to describe alternating patterning in the chick inner ear and has also been suggested to operate in the OoC in mammals via interactions of Nectins^21,22^. To test whether such a model captures the observed patterning features, we performed simulations where we assumed higher tension in SC:SC junctions compared to HC:SC junctions (Fig. S4a, Supplementary Video 10). The analysis showed that both the number of SC neighbors and the goodness of fit to hexagons are significantly higher compared to simulations that include global forces (Fig. S4b). Finally, we also checked how sensitive our model is to the choice of model parameters. We found that changes by ±50% of most parameters had a relatively modest effect on the values of the order parameters in the simulations (Fig. S4c).

As a mean to verify the model we sought to generate a prediction that can be tested experimentally by perturbing tissue mechanics. We focused on the effects of reducing the tensile forces in the tissue. Our model predicted that if the tension on all junctions was reduced and equated, the SC areas will increase relative to HC areas (Fig. 4a-b, Supplementary Video 11). To test this prediction experimentally, we added blebbistatin, a non-muscle myosin II (NMII) inhibitor, to E17.5 inner ear explants, 3 hours after the beginning of a time-lapse movie. It has been previously shown that adding blebbistatin to inner ear explants affects tension of SC:SC junctions and reduces movement of HCs ^7,23^. As predicted by our model, the addition of blebbistatin indeed cause a significant increase in SC areas compared to HC areas (Fig. 4c-d, Supplementary Video 12). Control explants where blebbistatin was not added did not show significant changes in HC and SC areas (Fig. 4d). These results support the assumption that an increase in SC:SC tension drives the decrease in SC areas and the corresponding increase in HC areas.

**Figure 4.**
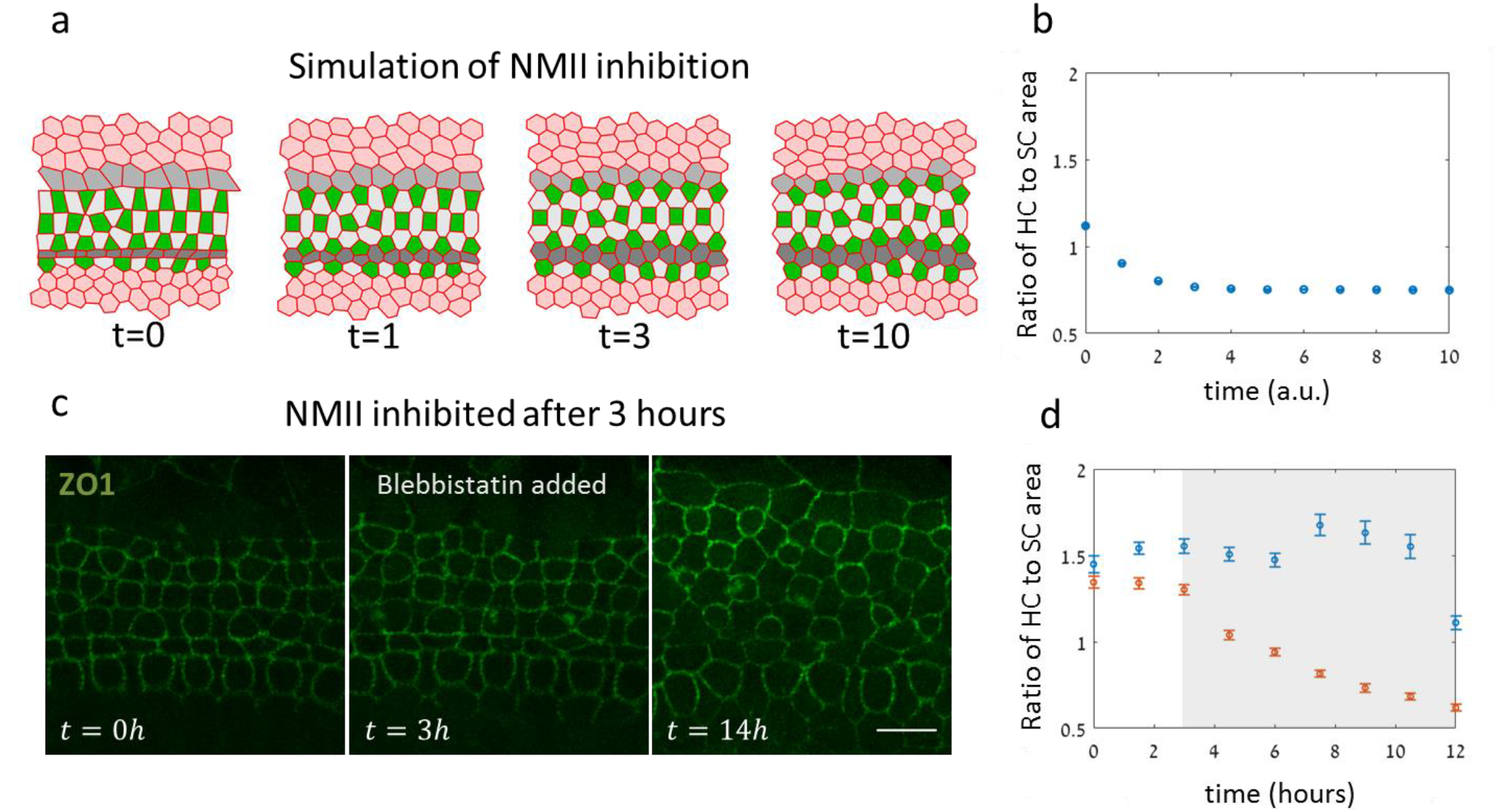
Model successfully predicts morphological changes induced by NMII inhibition. **(a)** A filmstrip of a simulation modeling NMII inhibition. The addition of a NMII inhibitor, blebbistatin, was simulated by reducing and equating the tension for all junction types and overall contractility. Simulation begins at the end point of the simulations shown in Fig. 3. Movie shown in Supplementary Video 11. **(b)** Analysis of the ratio of HC area to SC area during the simulations. Model predicts a decrease in the ratio of HCs relative to SCs. Data is an averaged over n=30 simulations. **(c)** A filmstrip from an E17.5 ZO1-GFP explant treated with blebbistatin (10 μM). Blebbistatin was added 3hrs after movie start. An increase in SCs area relative to HC area was observed within a few hours after addition of blebbistatin. Movie shown in Supplementary Video 12. Scale bar: 10 μm. **(d)** Analysis of the ratio of HC area to SC area as a function of time demonstrated a decrease in the ration of HC to SC areas in experiments where blebbistatin was added (red) compared to control experiments with no addition of blebbistatin. Data shows an average over n=3 experiments. Error bars in (b) and (d) indicate S.E.M.

Overall, our quantitative imaging analysis and mathematical modeling suggests that the formation of precise checkerboard-like pattern of OHCs is due to the combination of global shear and local repulsion forces between HCs. This mechanism is remarkably similar to crystal formation of particles (such as grains, atoms, or molecules) under global external force and local repulsion^24^. Likely sources for the shear and repulsion forces are the movement and division of Hensen cells, and steric repulsion between the nuclei of HCs, respectively. We also note that such shear forces are a potential mechanism for generating the spiral shape of the cochlea. Finally, we show that differential tension on SC:SC vs SC:HC junctions, regulated by NMII, drives the refinement of the OHC patterning. More generally, our proposed force-driven rearrangement mechanism may serve as a general principle for precise developmental patterning processes in other systems.

## Acknowledgement

We would like to thank members of the Sprinzak and Avraham labs for their advice and comments on this work. We would like to acknowledge Assaf Zaritsky for advice on image analysis and Dmitri Rivkin for help with image analysis. We would like to thank Steve Blacklow, Avigdor Eldar and Yasmine Meroz for fruitful discussions. Math1-EGFP mice were a gift from Jane E. Johnson, UT Southwestern.

## Contributions

This scientific study was conceived and planned by R.C., L.A.Z., and D.S. The inner ear explant experiments were performed by L.A.Z., R.C., S.W., and S.T. Support and expert advice for inner ear experiments were provided by K.B.A. The image analysis and image quantification were performed by R.C. and S.W., The ZO1- EGFP mouse and advice regarding its use were provided by F.M., The modeling was performed by R.C., M.H., S.B., and D.S. The manuscript was written by R.C., L.A.Z. and D.S.

## Methods

### Mice

The Math1-GFP line, J2XnGFP(Math1-nGFP tg) mice, were a gift from Jane E. Johnson, UT Southwestern^9^ and maintained on a C57BL/ 6 background. *Rosa26-ZO1-EGFP* mice were obtained from RIKEN Laboratory^13^ (accession no. CDB0260K; http://www2.clst.riken.jp/arg/reporter_mice.html) and maintained on a C57BL/6 background. All animal procedures were approved by the Animal Care and Use Committee at Tel Aviv University (04-16-014). Genotyping was performed using the KAPA HotStart Mouse Genotyping Kit (Sigma, KK7352) using GFP primers: Forward: TCCTTGAAGAAGATGGTGCG, Reverse: AAGTTCATCTGCACCACCG.

### Immunohistochemistry

Mice were sacrificed according to ethical guidelines and inner ears were dissected out in cold PBS and fixed in 4% paraformaldehyde (Electron Microscopy Sciences, cat: 15710) for 2 hrs at room temperature. Sensory epithelia were exposed and incubated in 1% normal Donkey serum (Sigma, cat: D9663) with 0.2% Triton-X (Sigma, cat: T-8787) for 2 hrs at room temperature. Samples were incubated with the ZO-1 primary antibody diluted 1:250 (Thermo Fisher Scientific, cat: 339100). Samples were incubated with secondary antibodies of Cy™3 AffiniPure Goat Anti-Mouse IgG (H+L) (Jackson laboratories, cat: 115-165-062). Stained samples were mounted on Cover glass 24×60mm thick. #1(0.13-0.17mm) slides (Bar-Naor Ltd. cat: BN1052441C) using a fluorescent mounting medium (GBI, cat: E18-18). Image acquisition was done with a ZEISS LSM 880 with Airyscan microscope (Zeiss). All images were taken from the apical area of the cochlea.

### Organ of Corti explants

The cochlea was dissected using sterile conditions under Nikon SM2 745T Stereomicroscope (Nikon). Explant cultures were prepared from the cochleae of *ZO1-GFP* transgenic mice. The cochleae were removed and placed in ice cold PBS. Using two forceps the organ of Corti was gently freed from the capsule and separated from the stria vascularis. The tissue was oriented so that the apical surfaces of the hair cells were pointing down, directed toward the Matrigel Phenol Red Free solution (In vitro technologies, cat: FAL356237). Excess medium was removed and explanted tissue was allowed to attach to the Matrigel for 5-10 min in a 37°C incubator with 5% CO2 while avoiding drying of the tissue. After tissue attachment, Dulbecco’s modified Eagle’s medium (Biological industries, cat: 01-053-1A) supplemented with N-2 Supplement (100X) (Thermo Fisher Scientific, cat: 17502001) and 1% FBS (Biological industries, cat: 04-007-1A) was added gently. The plate was then placed in the 37°C incubator of the microscope and as a control in the lab incubator. Cultures were kept for up to 48 hrs.

### Microscopy details

Cochleae were imaged using Zeiss LSM 880 confocal microscope equipped with an Airyscan detector using 488nm laser for GFP and 561nm laser for Cy3. For fixed samples we used Plan-apochromat 63× oil-immersion objective with NA=1.4. For live imaging we used C-apochromat 40× water-immersion objective with NA=1.2. The microscope was equipped with a 37 °C temperature-controlled chamber and a CO2 regulator providing 5% CO2. The equipment was controlled using Zeiss software – “Zen black”.

### Image analysis

Images of fixed samples were taken at high resolution (63× objective) and tiled together to form a full image of the cochlea. For fixed samples we used cochleae from transgenic mice expressing Math1-GFP, which is an early marker for HCs. Since the apical side of the OoC is usually tilted relative to the focal plane, 15-30 z-stacks separated by 1μm were taken at each position for both fixed and live imaging. The stacks were then projected to the focal plane using a maximum intensity projection method.

All data processing was performed off-line using custom built code in Matlab (MATLAB R2018b, the MathWorks Inc.). A semi-automatic analysis code was used for segmentation of the boundaries of the cells and for data extraction. In short, segmentation was done by applying filters to the image and using a watershed algorithm. Segmentation defects were corrected manually using a custom-made Matlab GUI. Different cell types for OHC (OHC1 or OHC3), OHC2 or SC were manually marked relying on the Math1-GFP marker. In early stages of development, where the HC rows are not yet formed, we define OHC2 as a HC that neither borders the PCs row nor the lateral border of the OoC.

The initial segmentation output provides information about the identity (cell type), position (centroid coordinates), shape (area) and connectivity (number of neighbors, contact lengths) of each cell. We focus on analysis of the middle row of OHCs, OHC_2_, since the HCs in this row only border SCs within the OHC region (i.e. in the final pattern they should have exactly 4 SC neighbors). The morphological and order parameters presented in Fig. 1e-g are evaluated for each per cell in OHC2 using the segmentation output. These parameters are then grouped by the cochlear positions of the cells (see insert in Fig. 1e) and averaged. The morphological parameters and number of SC neighbors are directly extracted from the segmentation output.

The goodness of fit to hexagon was calculated for each HC in OHC2. The general idea is to find how well do the centroids of the neighboring HCs to the cell of interest (COI) fits to a symmetric hexagon (defined below). Neighboring HCs of a COI are defined as cells with a shared junction in the Voronoi tessellation made by the centroids of all the HCs. The reason that we fit elongated hexagons and not regular hexagons is that sometimes the apical surface of the OoC have some relative angle with respect to the focal plane, so that the projected analyzed image can be slightly “squeezed”.

Once we identified the neighboring HCs to the COI, we search for symmetric hexagon with vertices that fits best to the centroids of the neighboring HCs. We define a symmetric hexagon by starting with a regular hexagon with vertices at:

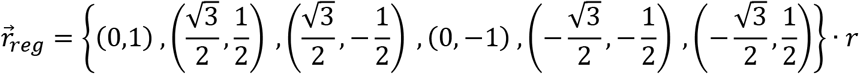

where *r* defines the size of the hexagon. We then introduce an additional degree of freedom for scaling in the *x* axis (effectively creating an elongated regular hexagon):

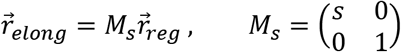

where *s* is the scaling factor. Since the general orientation of the symmetric hexagon can be different at different positions along the cochlea, we allow another degree of freedom in the form of a simple rotation:

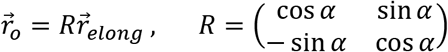

where α is the rotation angle. Finally, the symmetric hexagon should be centered at the centroid of the COI, so the fitted hexagon vertices are:

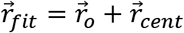

Where 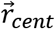 is the centroid of the COI. The best fit to symmetric hexagon is defined as the minimal sum of weighted squared differences between the centroids of neighboring HCs and 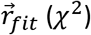 The value of *χ*^2^ is normalized by the number of independent degrees of freedom (# HC neighbors minus the # of fit parameters). The fit parameters are *r*, *s* and α. The weights are taken to be equal to *r*^−1^.

#### Pixel displacement analysis

To analyze the shear profile observed in movies along the base-to-apex axis we rotate the movies so that the IHCs are at the bottom of the field of view and the OHCs are at the top. In addition to the relative motion between the cells, there is a global motion of the tissue. To consider only relative motion, we set the PC row to be static with respect to the field of view, using image registration algorithm (Matlab – “imregdemons”). Then, using the same algorithm, for each pixel we sum the pair-wise translations between pairs of frames to get the total displacement of the pixel at the end of the movie. The analysis showed that pixels initially located higher in the field of view had larger total displacement relative to pixels that started at lower position, as expected from a shear motion profile.

#### Intercalations analysis

This analysis was done manually with the aid of an assisting GUI. The intercalations were identified visually and counted by clicking the location of intercalation. Each of these positions were marked for the duration of the intercalation to prevent repeated counts of the same transition. The intercalations were counted only for the OHC region as the IHC region is mostly organized at the developmental times we investigate. Finally, the total number of intercalations was divided by the number of cells in the OHC region and the duration of the movie.

### Cellular area movie analysis

Movies of samples treated with blebbistatin were analyzed in the same way as fixed samples. The borders of the cells were segmented and the areas of the cells together with their identity were extracted from the segmentation output. For each analyzed frame in the movie, the average surface areas of HCs and SCs were calculated and then divided to get the ratio of HC to SC area. As a control, the same analysis was done for movies with no addition of blebbistatin.

#### Calculation of correlations and their p-value

All statistical analysis was performed using Pearson correlation. The number of samples is as indicated in each figure caption.

### Details of the model

#### Mechanical energy function for the 2D vertex model

To describe the dynamics of the apical surface of the OoC we use 2D vertex model previously used to describe morphological processes of epithelial tissues ^15,16^. In this model each cell is described as a polygon which is uniquely defined by the position of its edges and vertices. A total mechanical energy associated with the position of the vertices, length of edges, and areas of the cells is minimized at each time step.

The energy function (also shown in Fig. S3) is given by:

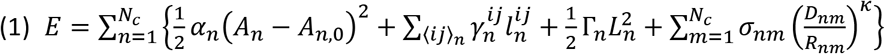

where *n*, *m* are cell indices, *N_c_* is the total number of cells and 〈*ij*〉_*n*_ are the pairs of adjacent vertices in cell *n* (between vertices *i* and *j*).

The first element in the energy function is an elastic term in the area of the cell *A*_*n*_; each cell has a preferable area *A*_0,*n*_ and each deviation from it results in a quadratic energy cost. Similar to a spring constant describing the rigidity of a spring, α_*n*_ describes the incompressibility of cell *n*. The second element is a linear term in the length of the junction *l*_*i*,*j*_. The parameter *γ*_*ij*_ describes the tension of junction *ij* in the sense that any increase in the junction’s length results in an energy cost directly proportional to *γ*_*ij*_. The third element in a quadratic term in the perimeter *L*_*n*_ of cell *n,* and represents the susceptibility of the cells to deformations. Γ_*n*_ describes the contractility of the cell, or the tendency of the cell to contract, and hence the tendency of the cell to round up (physically controlled by actomyosin cables in the cell cortex).

#### Local repulsion between HCs

In addition to these standard terms in the energy function, we also define additional terms associated with global shear and local repulsion. Since some of these forces are affecting only HCs and not SCs we now associate each cell with a different identity in the OoC. Overall we define the following cell types: HC, SC, PC, top boundary cell or general cell outside the OoC. This also allows setting different mechanical parameters for different cell types and junction types.

The local repulsion is set in the energy function by the last term in eq. 1. As mentioned in the text, we assume that the repulsion forces can result of interactions between HCs at nuclei level. This is described by an effective hard sphere interaction between the HCs. The distance between the pair of cells *n* and *m*, *R*_*nm*_, is defined as the distance between the centers of masses of the cells. *σ*_*nm*_, *D*_*nm*_ and *κ* quantifies the strength and behavior of these repulsion forces. Taking a high value for *κ* results in a smooth version of a hard-sphere potential with a radius *D*_*nm*_/2. Since the repulsion forces act only between HCs, we effectively have:

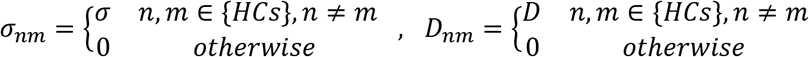

#### Global shear forces in the model

To model the shear force observed in experiments we decompose this force to two components of horizontal shear and vertical pull. Horizontal shear is characterized by a velocity gradient vertical to the direction of motion, namely, if the motion is in the *x* direction, the velocity changes with *y*. Hence, we can take this component to be proportional to the *y* coordinate of each cell. For the vertical pull component we assume a quadratic energy term with respect to the *y*, namely, 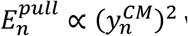 where 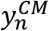 is the *y* coordinate of the center of mass of cell *n* from the PCs. Hence, the pull force is proportional to the distance of the cell from the PCs. Overall, the global shear forces are taken into account by assuming external forces that acts on vertex *i* of cell *n* given by:

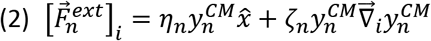

where *η*_*n*_ and *ζ*_*n*_ describe the horizontal shear and vertical pull that act on HCs and 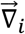 is the divergence relative to vertex *i* position. Since the shear and pulling forces act only on HCs, we effectively have

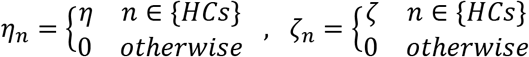

The parameters values for different cell types and different junction types are given in table S1.

To account for random fluctuations of the vertices, we also add random noise for each of the vertices components as 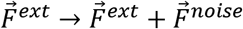 and randomize the components of 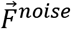 every few timesteps.

#### Initial and boundary conditions

To generate initial disordered 2D lattices, we use the method described in ^25^. In short, we begin with 12×12 hexagonal lattices and then run the 2D vertex simulations with assigning random values for the tension (*γ*_*i*,*j*_) and preferable areas (*A*_0,*n*_). For simplicity, we use periodic boundary conditions at the edges of the cell lattice.

As mentioned in the main text, we focus on the organization of the OHC region, so we begin the simulations in an initial state where the IHC and PC rows already formed. The PCs form straight boundaries both with the IHC region and with the OHC region (Fig. 1c,d). In the simulation this is achieved by setting high tension (*γ*_*i*,*j*_) in junctions that separate PCs from non-PCs. Since the tension in the boundary junctions is higher, the model minimizes the boundary length to a straight line. In the rest of the simulation the PC row serves as a boundary condition for the patterning of the outer HCs. To simulate the differentiation pattern of OHCs into HCs and SCs via lateral inhibition, we first define a region above the pillar cells in which lateral inhibition is active. We then apply the lateral inhibition process^17-19^ by picking a random cell from this region and letting it differentiate into a HC only if it has no HC neighbors. This process is repeated until no additional hair cells can differentiate (according to the lateral inhibition rules). We note that we allow the differentiation of new HCs throughout the simulations according to the same rules, namely, if there is a SC within the OHC region that does not touch a HC, that cell can differentiate into a HC.

#### Running the simulations

The 2D vertex model simulates the dynamics of epithelial tissues by minimizing the net force on the system. Each term in the energy function and force expression can be calculated from the positions of all the vertices (given a set of model parameters). In other words, the energy is a function of the position of all the vertices. For the purpose of convenience, the position of all the vertices is defined through a single vector structured such that entries (2*i* − 1, 2*i*) of the vector represents the (*x*, *y*) position of vertex *i* respectively. If the total number of vertices is *N*_*v*_, the position vector looks like this:

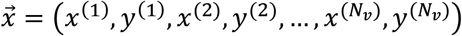

where *x*^(*i*)^, *y*^(*i*)^ are the (*x*, *y*) positions of vertex *i*. We assume that at each moment the system is trying to reach a steady state of zero net force on all of the vertices, meaning:

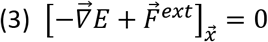

where 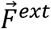 is a vector of size *N*_*v*_ with the external force components of the vertices in the same format as 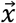 and the gradient is defined as: 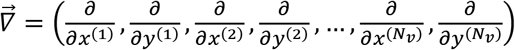

We can iteratively advance the vertices position according to the gradient descent method in the following way:

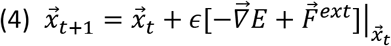

where *t* indicates the time step and *ϵ* is a small enough iteration parameter.

In parallel to advancing to the mechanical steady state, we introduce T1 and T2 transitions. A T1 transition initiates for junctions smaller than some threshold length with some probability. A T2 transition will initiate for each cell with area smaller than some threshold area.

Once T1 transition has occurred, the initial transitioned junction is small and may go through a T1 process again. To prevent a situation where a certain junction goes through intercalations over and over at the same point, we allow a short relaxation time for the transitioned junction when intercalations are not allowed.

Finally, to get exactly three rows of OHCs in the simulations, we limit the number of cells differentiating into OHCs so that the cells fit into exactly three rows (without extra HCs in the 4^th^ row). We note, that this is necessary because of the imposed periodic boundary conditions in the simulations. Without the periodic boundary conditions, extra OHCs can be pushed forward towards the apex making sure an exact number of OHCs that fits three rows is reached.

All simulation codes were uploaded to public repository https://github.com/Roie-Cohen/Inner_ear_vertex_code.

#### Extracting statistics from the simulations

For each time step in each simulation, statistics were taken in the same manner as in the analysis of fixed samples. The statistics include the number of SC neighbors, the goodness of fit to hexagon and the HC area. Statistics are taken for HCs that don’t touch the PCs or the top border. For Stage 1, we introduce the top border and only then take the statistics. For each time step, the statistics were averaged over all the simulations.

One difference between the real tissue and the model is the lack of curvature of the junctions in the simulations. At the end of development HCs are circular and close packed, and this gives drastically different surface area of HCs in the model relative to the real tissue. To account for this, for each of the HCs we find the closest HC neighbor and define its radius as half the distance between the two cells. The effective area for each HC is defined as the area of the representing circle. For other cell types (such as SCs), the overlapped area between the circular HCs and the cell of interest is reduced from the polygonal area of the cell.

#### Modeling blebbistatin addition and alternative models

In simulations modeling blebbistatin treatment, we begin at the last step of Stage 2 and then reduce and equate the tension by setting 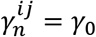 for all junction types. Additionally, shear and compression are suppressed by taking *η* = ζ = 0. Parameters for these simulations are shown in table S1.

In the alternative model, the assumption is that the organization forms only due to tensile differences between different cellular junctions. More specifically, we set *η* = *ζ* =0 and there is no difference in compressibility and contractility between HCs and SCs (*α*_*n*_ = *α*_0_, Γ_*n*_= Γ_0_ for all *n*). Parameters for these simulations are shown in table S1.

#### Model parameters

The parameter values used in the simulations are detailed in table S1.

## Appendix calculation of center of mass derivatives

Although all of the variables in eq. 1-2 are a function of the vertices position, the center of mass of a cell does not have a trivial form. In order to calculate the center of mass of the cells in the model, we use the formula for the center of mass of a n-sided polygon. Given an ordered set of vertices position {*x*_*i*_, *y*_*i*_} with *i* = 0,1 … *n* − 1, the center of mass coordinated are:

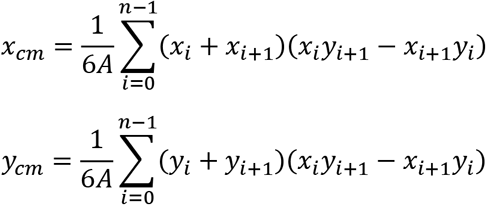

where the area of the polygon *A* is given by:

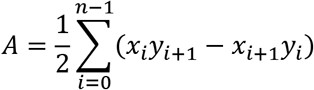

In these formulas we define *x*_*n*_ ≡ *x*_0_ and *y*_*n*_ ≡ *y*_0_. As part of the derivatives in eq. 3, we use the derivatives of the center of mass relative to the positions of the vertices. For one of the vertices, *x*_*k*_, *y*_*k*_these derivatives are given by:

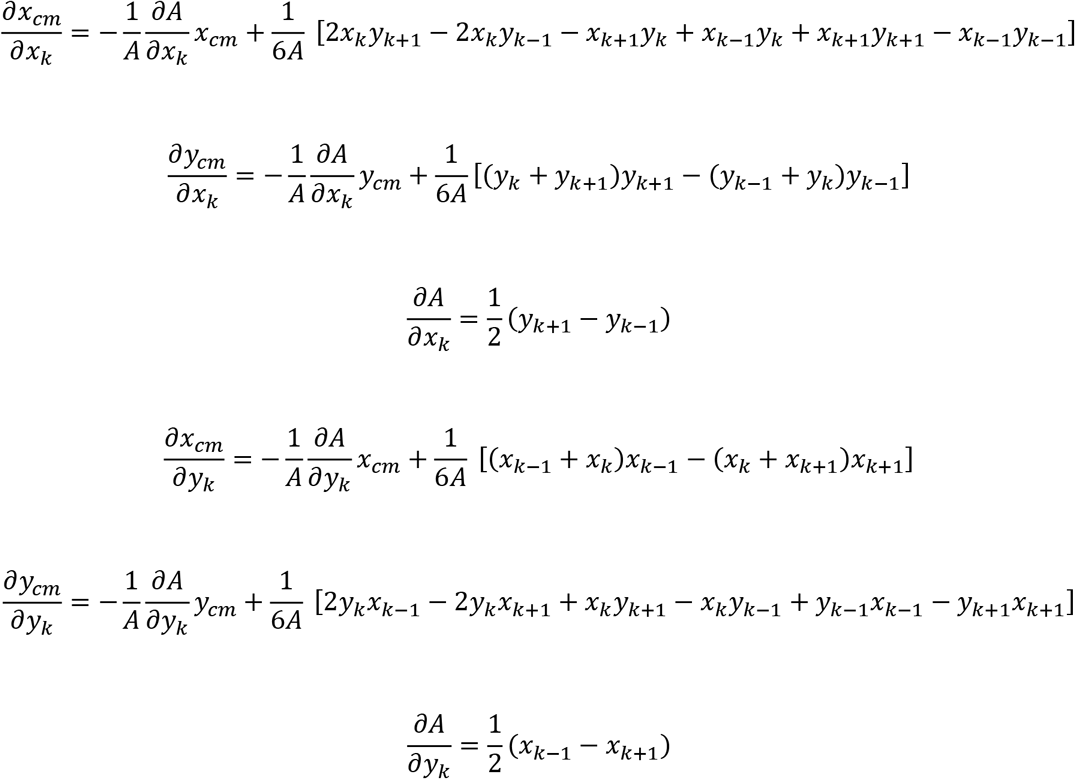

In the repulsive term in eq. 1, the enter of mass is hidden in *R*_*nm*_:

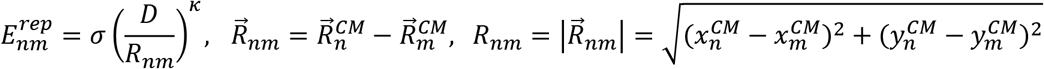

where 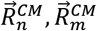 are the center of mass position of the cells. For a vertex *k* of cell *n*, the derivative of this energy term with respect to the position of vertex *k* is calculated as:

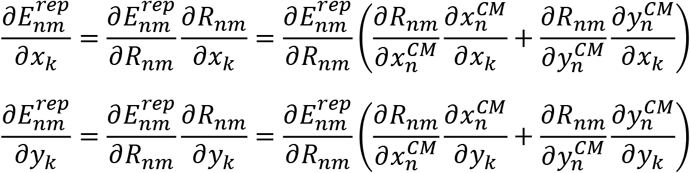

---

**Figure S1.**
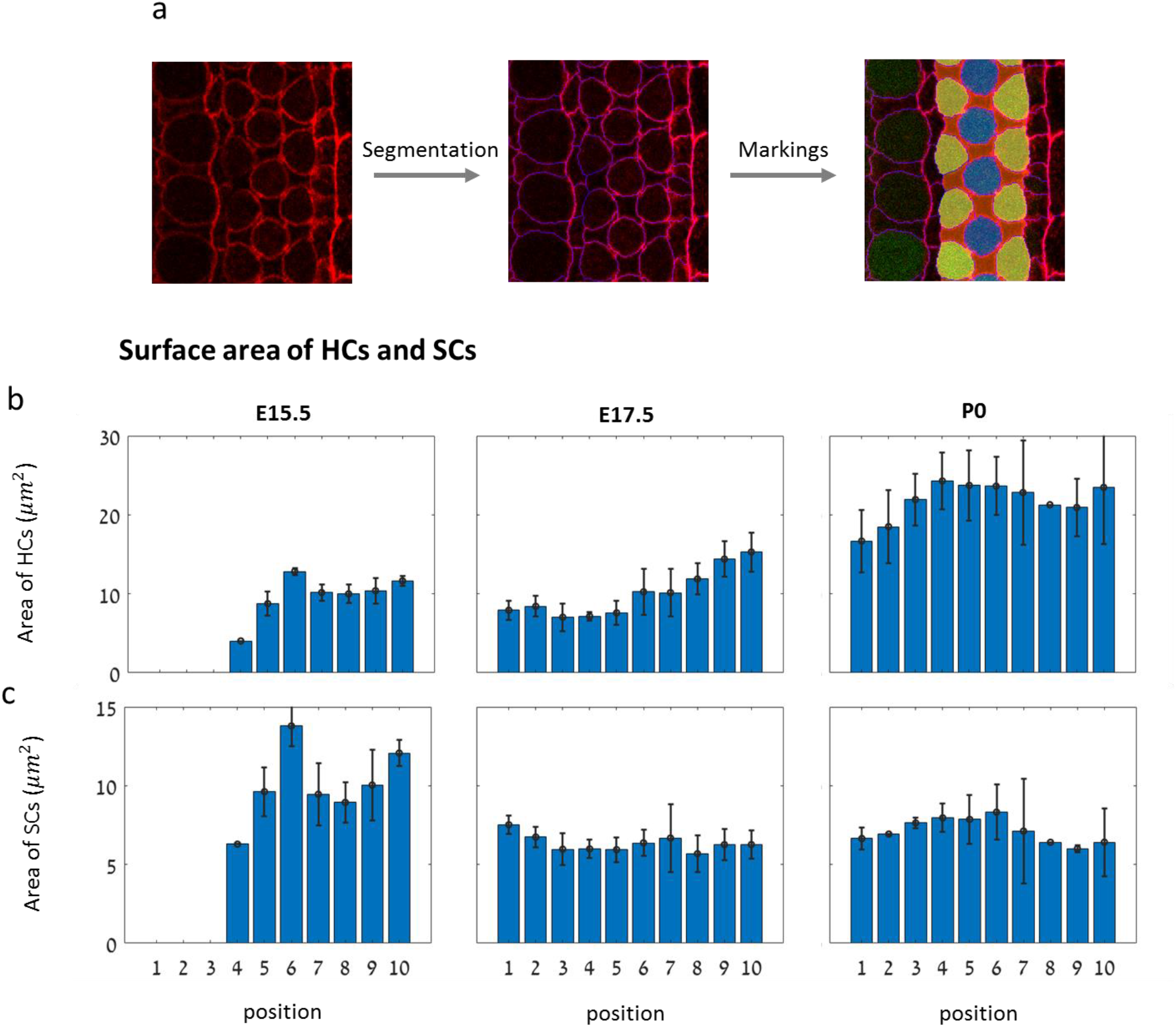
Description of image analysis sequence and additional morphological data on HCs and SCs. **(a)** Schematic of the segmentation process. The borders of cells are delineated (purple lines) and different cell types are marked (yellow/orange/blue cells) using a semi-automatic code. **(b-c)** Surface area of HCs (b) and SCs (c) in different regions of the cochlea from apex to base at E15.5, E17.5, P0. This analysis shows that as development progresses, HC surface area increases while SC surface area slightly decreases. The data is averaged over n=3,4,3 cochleae at E15.5, E17.5, P0, respectively. Error bars indicate S.E.M

**Figure S2.**
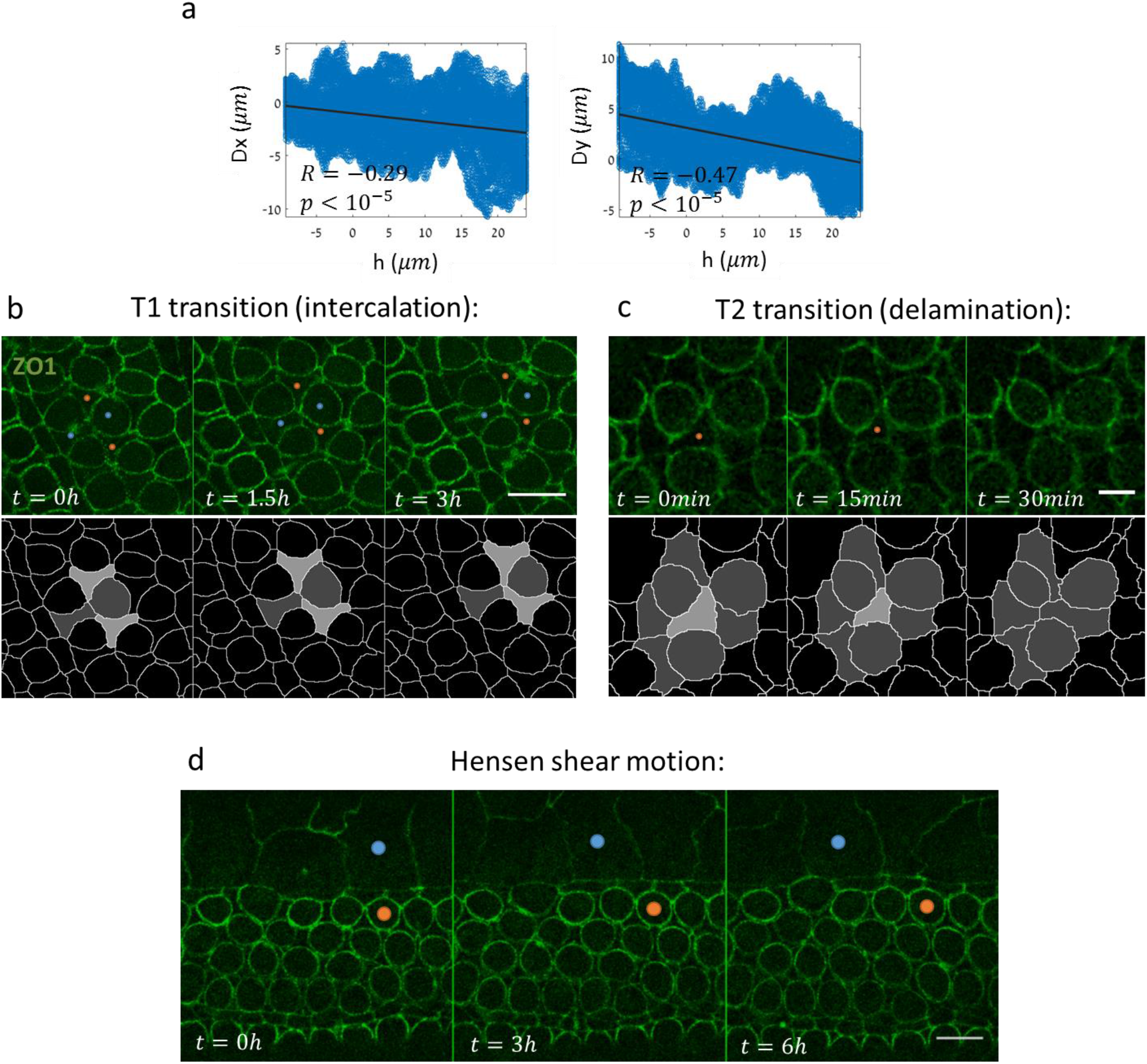
Additional data on shear movement, intercalation and delaminations. **(a)** Total pixel displacement as a function of the initial distance from the PCs acquired from another time-lapse movie of an E15.5 cochlear explant (different sample than Fig. 2a-b). Black lines represent linear regressions for the horizontal (left) and perpendicular (right) movements. The correlation coefficients (R) and corresponding p-values are as denoted. **(b)** A filmstrip from a E17.5 mouse cochlear explant showing an intercalation process (T1 transition) between the cell pair marked in red dots and the cell pair marked in blue dots. Bottom row shows a segmented version of the intercalation process. Movie shown in Supplementary Video 5. Scale bar: 10*μm*. **(c)** A filmstrip from a E17.5 mouse cochlear explant showing a delamination process (T2 transition), where the cell marked with red dot decreases in surface area and eventually vanishes from the apical surface. Bottom row presents a segmented version of the transition. Movie shown in Supplementary Video 6. Scale bar: 5*μm*. **(d)** Filmstrip showing sliding motion of Hensen cells on E17.5 cochlear explant. Blue and red dots mark initially aligned Hensen cell and OHC, respectively. Movie shown in Supplementary Video 2. Scale bar: 10*μm*

**Figure S3.**
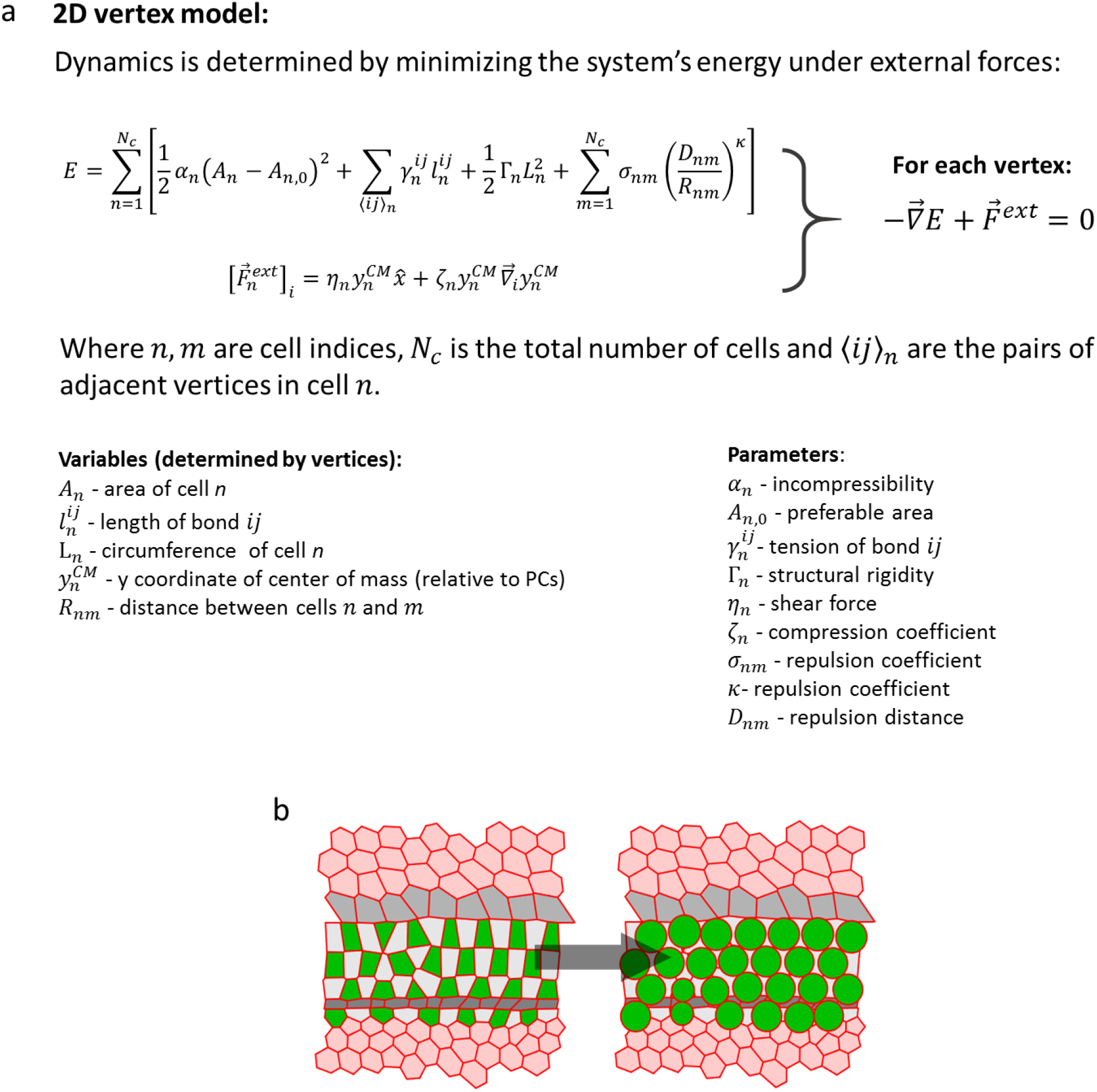
(a) Mathematical description of the model. The dynamics of the vertices (e.g. their position) is determined by minimizing the mechanical energy function of the system under external forces. The energy function includes terms corresponding to internal pressure (depends on cell area), junction tension (depends on junction length), cell contractility (depends on cell perimeter), and repulsion forces between HCs (depend on distance between HCS). In addition to the energy function there are external forces (non-conserved) that correspond to horizontal shear and vertical compression (both depend on distance from PCs). Bottom: list of variables (depend on vertex positions) and parameters of the model. **(b)** Schematic of the process of HC rounding. Each HC was replaced by a circle whose radius is half the distance to its closest HC neighbor.

**Figure S4.**
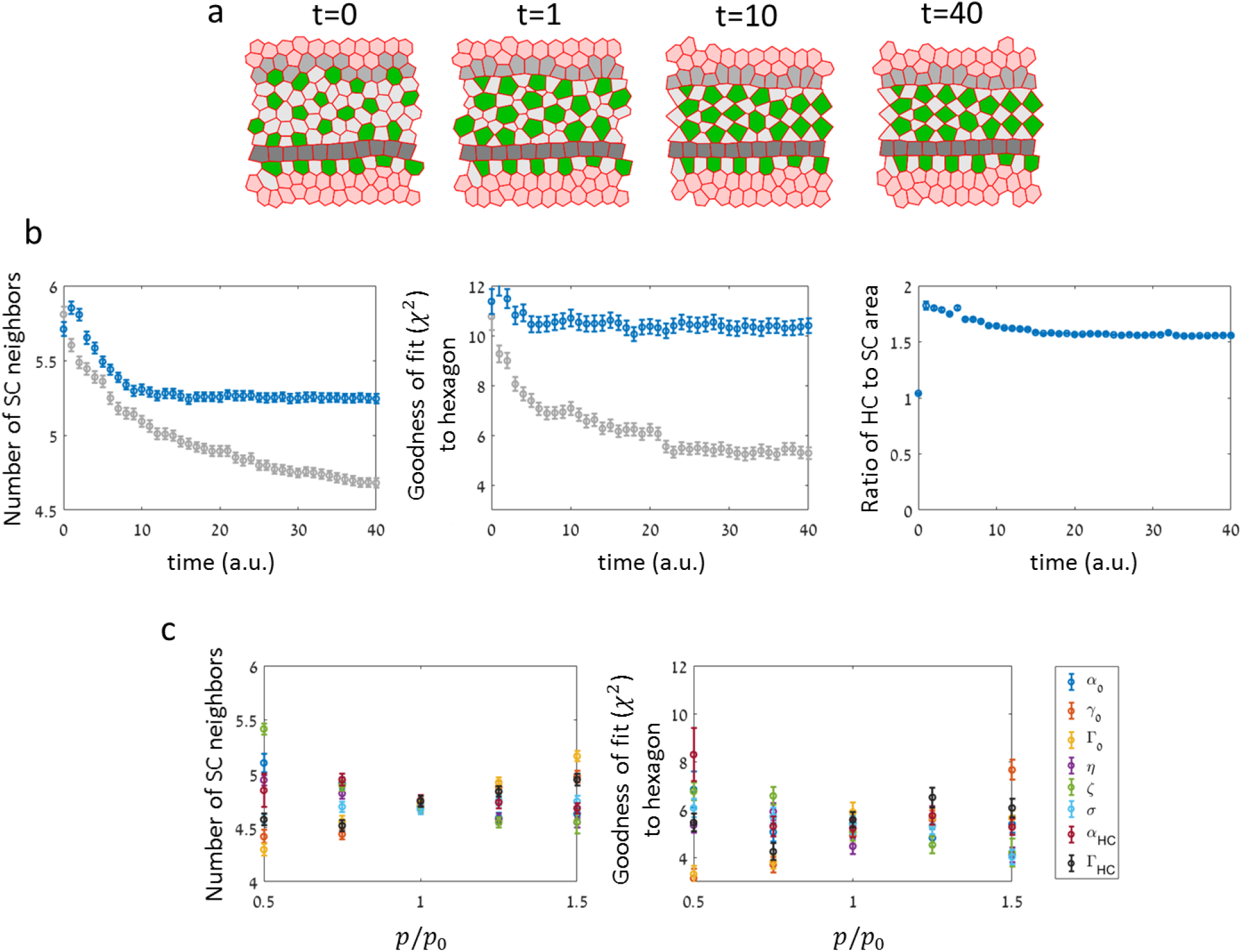
Alternative models and robustness to parameters. **(a)** A filmstrip of a simulation from an alternative model that do not include global shear and local repulsion. The alternative model consists only of the second stage in the original model, meaning higher tension in SC:SC junctions relative to HC:SC junctions. Movie shown in Supplementary Video 10. **(b)** Simulations of the alternative model failed to capture the transition to organized pattern compared to the original model. Although the number of SC neighbors decreased and the HC organization became more hexagonal (blue markers), simulation from the original model (grey markers) produced order parameters closer to those observed experimentally. The data was averaged over n=50 simulations. Error bars indicate S.E.M. Full description of the simulations is provided in the methods. Parameters used are provided in table S1. **(c)** Parameter sensitivity. Order parameters produced by running simulations with different values of model parameters (in the original model). The parameters were evaluated at the end of the 1^st^ stage of the original model. *p*/*p*_0_ represents a normalized value of the model parameter, where *p*_0_ is the value of the parameter in the original model. Legend indicate parameters used in the analysis. This data shows that changes of ±50% in the model parameters produced relatively small changes in the values of order parameters, showing that the model is robust to model parameter values.

**Figure S1.**
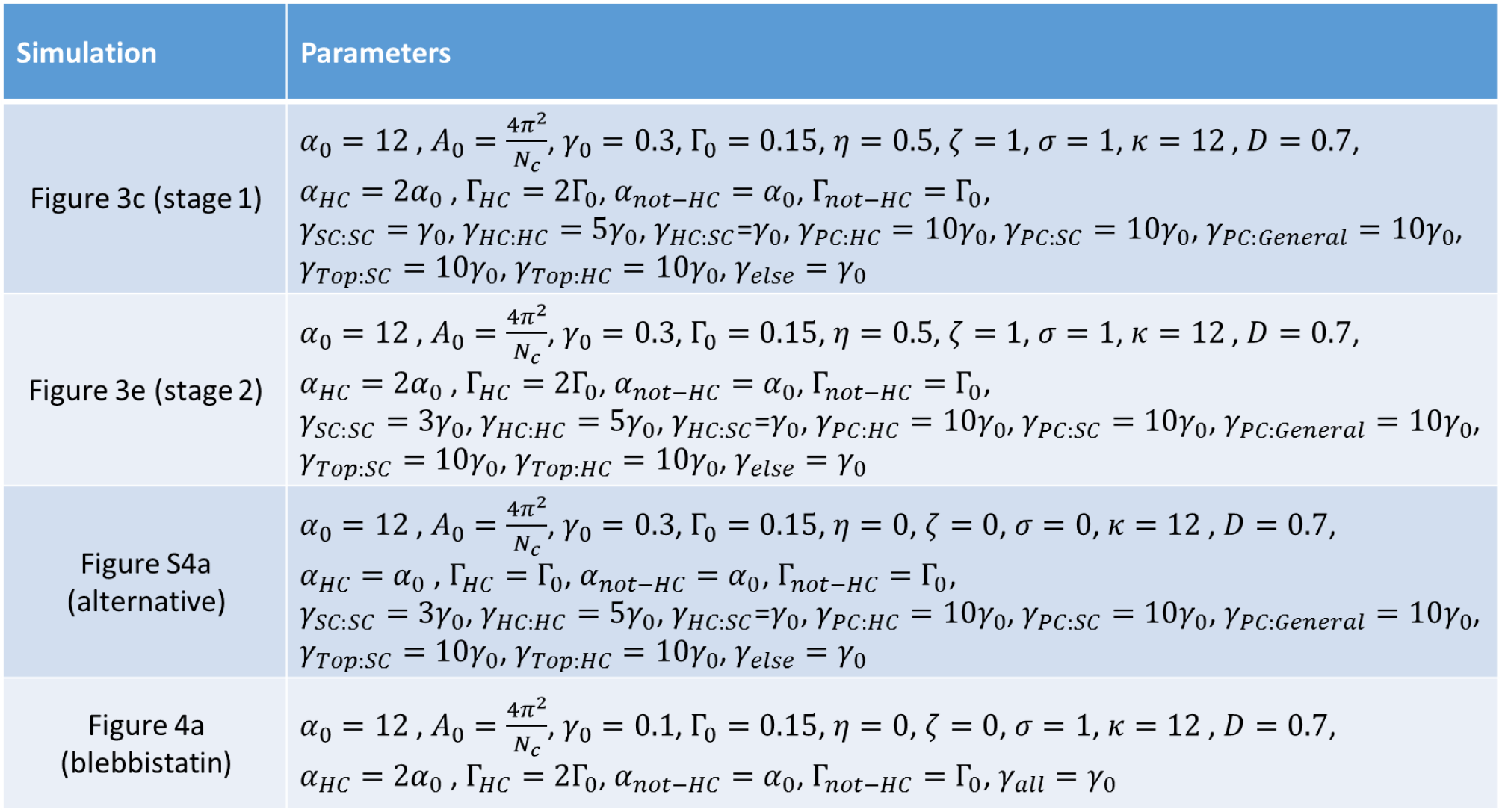
Model parameters used in simulations.

## Supplementary videos

**Supplementary Video 1.** A movie from the apex region of a cochlear explant of a ZO1-GFP mouse at E15.5, showing shear motion along the base-to-apex direction. Movie corresponds to the filmstrip shown in Fig. 2b. Scale bar: 10*μm*.

**Supplementary Video 2.** A movie of a cochlear explant of a ZO1-GFP mouse at E17.5, showing shear motion of the Hensen cells along the base-to-apex direction. Movie corresponds to the filmstrip shown in Fig. S2d. Scale bar: 10*μm*.

**Supplementary Video 3.** A movie from the apex region of a cochlear explant of a ZO1-GFP mouse at E15.5, showing an intercalation process (T1 transition) between the cell pair marked in red dots and the cell pair marked in blue dots. Movie corresponds to the filmstrip shown in Fig. 2d. Scale bar: 5*μm*.

**Supplementary Video 4.** A movie from the apex region of a cochlear explant of a ZO1-GFP mouse at E15.5, showing a delamination process (T2 transition), where the marked cell decreases in surface area and eventually vanishes from the apical surface. Movie corresponds to the filmstrip shown in Fig. 2e. Scale bar: 5*μm*.

**Supplementary Video 5.** A movie from a E17.5 mouse cochlear explant showing an intercalation process (T1 transition) between the cell pair marked in red dots and the cell pair marked in blue dots. Movie corresponds to the filmstrip shown in Fig. S2b. Scale bar: 5 *μm*.

**Supplementary Video 6.** A movie from a E17.5 mouse cochlear explant showing a delamination process (T2 transition), where the cell marked with red dot decreases in surface area and eventually vanishes from the apical surface. Movie corresponds to the filmstrip shown in Fig. S2c. Scale bar: 5 *μm*.

**Supplementary Video 7.** A movie from the apex region of a cochlear explant of a ZO1-GFP mouse at E15.5, showing an event where a cell (marked) is “squeezed out” towards the top border of OHC region. Movie corresponds to the filmstrip shown in Fig. 2g. Scale bar: 5*μm*.

**Supplementary Video 8.** A movie of the 1^st^ stage in the simulation. Global shear and local repulsion forces lead to the formation of a compact state of HCs. Movie corresponds to the filmstrip shown in Fig. 3c.

**Supplementary Video 9.** A movie of the 2^nd^ stage in the simulation. This stage begins at the end of the compaction stage (end of Supplementary Video 8). Movie corresponds to the filmstrip shown in Fig. 3e.

**Supplementary Video 10.** A movie of a simulation from the alternative model. The alternative model consists only of the 2^nd^ stage in the original model, meaning higher tension in SC:SC junctions relative to HC:SC junctions. Movie corresponds to the filmstrip shown in Fig. S4a.

**Supplementary Video 11.** A movie of a simulation modeling NMII inhibition. The addition of a NMII inhibitor, blebbistatin, is simulated by reducing and equating the tension for all junction types and overall contractility. Simulation begins at the end point of the simulations shown in Fig. 3. Movie corresponds to the filmstrip shown in Fig. 4a.

**Supplementary Video 12.** A movie from an E17.5 ZO1-GFP explant treated with blebbistatin (10 μM). Blebbistatin added 3hrs after movie starts. An increase in SCs area relative to HC area was observed within a few hours after addition of blebbistatin. Green dot in the upper right corner indicates the addition of blebbistatin. Movie corresponds to the filmstrip shown in Fig. 4c. Scale bar: 10 μm.

## References

1 Basch, M. L., Brown, R. M., 2nd, Jen, H. I. & Groves, A. K. Where hearing starts: the development of the mammalian cochlea. J Anat 228, 233–254, doi:10.1111/joa.12314 (2016).

2 Kelly, M. C. & Chen, P. Development of form and function in the mammalian cochlea. Curr Opin Neurobiol 19, 395–401, doi:10.1016/j.conb.2009.07.010 (2009).

3 Bryant, J., Goodyear, R. J. & Richardson, G. P. Sensory organ development in the inner ear: molecular and cellular mechanisms. Br Med Bull 63, 39–57, doi:10.1093/bmb/63.1.39 (2002).

4 Wang, J. et al. Regulation of polarized extension and planar cell polarity in the cochlea by the vertebrate PCP pathway. Nat Genet 37, 980–985, doi:10.1038/ng1622 (2005).

5 Yamamoto, N., Okano, T., Ma, X., Adelstein, R. S. & Kelley, M. W. Myosin II regulates extension, growth and patterning in the mammalian cochlear duct. Development 136, 1977–1986, doi:10.1242/dev.030718 (2009).

6 Chacon-Heszele, M. F., Ren, D., Reynolds, A. B., Chi, F. & Chen, P. Regulation of cochlear convergent extension by the vertebrate planar cell polarity pathway is dependent on p120-catenin. Development 139, 968–978, doi:10.1242/dev.065326 (2012).

7 Driver, E. C., Northrop, A. & Kelley, M. W. Cell migration, intercalation and growth regulate mammalian cochlear extension. Development 144, 3766–3776, doi:10.1242/dev.151761 (2017).

8 Lee, Y. S., Liu, F. & Segil, N. A morphogenetic wave of p27Kip1 transcription directs cell cycle exit during organ of Corti development. Development 133, 2817–2826, doi:10.1242/dev.02453 (2006).

9 Lumpkin, E. A. et al. Math1-driven GFP expression in the developing nervous system of transgenic mice. Gene Expr Patterns 3, 389–395 (2003).

10 Fanning, A. S. & Anderson, J. M. Zonula occludens-1 and -2 are cytosolic scaffolds that regulate the assembly of cellular junctions. Ann N Y Acad Sci 1165, 113–120, doi:10.1111/j.1749-6632.2009.04440.x (2009).

11 Dror, A. A. & Avraham, K. B. Hearing loss: mechanisms revealed by genetics and cell biology. Annu Rev Genet 43, 411–437, doi:10.1146/annurev-genet-102108-134135 (2009).

12 McKenzie, E., Krupin, A. & Kelley, M. W. Cellular growth and rearrangement during the development of the mammalian organ of Corti. Dev Dyn 229, 802–812, doi:10.1002/dvdy.10500 (2004).

13 Katsunuma, S. et al. Synergistic action of nectins and cadherins generates the mosaic cellular pattern of the olfactory epithelium. J Cell Biol 212, 561–575, doi:10.1083/jcb.201509020 (2016).

14 Marinari, E. et al. Live-cell delamination counterbalances epithelial growth to limit tissue overcrowding. Nature 484, 542–545, doi:10.1038/nature10984 (2012).

15 Farhadifar, R., Roper, J. C., Aigouy, B., Eaton, S. & Julicher, F. The influence of cell mechanics, cell-cell interactions, and proliferation on epithelial packing. Curr Biol 17, 2095–2104, doi:10.1016/j.cub.2007.11.049 (2007).

16 Hufnagel, L., Teleman, A. A., Rouault, H., Cohen, S. M. & Shraiman, B. I. On the mechanism of wing size determination in fly development. Proc Natl Acad Sci U S A 104, 3835–3840, doi:10.1073/pnas.0607134104 (2007).

17 Collier, J. R., Monk, N. A., Maini, P. K. & Lewis, J. H. Pattern formation by lateral inhibition with feedback: a mathematical model of delta-notch intercellular signalling. J Theor Biol 183, 429–446, doi:10.1006/jtbi.1996.0233 (1996).

18 Shaya, O. & Sprinzak, D. From Notch signaling to fine-grained patterning: Modeling meets experiments. Curr Opin Genet Dev 21, 732–739, doi:10.1016/j.gde.2011.07.007 (2011).

19 Sprinzak, D. et al. Cis-interactions between Notch and Delta generate mutually exclusive signalling states. Nature 465, 86–90, doi:10.1038/nature08959 (2010).

20 Chiou, K. K., Hufnagel, L. & Shraiman, B. I. Mechanical stress inference for two dimensional cell arrays. PLoS Comput Biol 8, e1002512, doi:10.1371/journal.pcbi.1002512 (2012).

21 Honda, H., Yamanaka, H. & Eguchi, G. Transformation of a polygonal cellular pattern during sexual maturation of the avian oviduct epithelium: computer simulation. J Embryol Exp Morphol 98, 1–19 (1986).

22 Togashi, H. et al. Nectins establish a checkerboard-like cellular pattern in the auditory epithelium. Science 333, 1144–1147, doi:10.1126/science.1208467 (2011).

23 Ebrahim, S. et al. NMII forms a contractile transcellular sarcomeric network to regulate apical cell junctions and tissue geometry. Curr Biol 23, 731–736, doi:10.1016/j.cub.2013.03.039 (2013).

24 Tsai, J. C., Voth, G. A. & Gollub, J. P. Internal granular dynamics, shear-induced crystallization, and compaction steps. Phys Rev Lett 91, 064301, doi:10.1103/PhysRevLett.91.064301 (2003).

25 Shaya, O. et al. Cell-Cell Contact Area Affects Notch Signaling and Notch-Dependent Patterning. Dev Cell 40, 505–511 e506, doi:10.1016/j.devcel.2017.02.009 (2017).

